# Laurdan discerns lipid membrane hydration and cholesterol content

**DOI:** 10.1101/2022.11.02.514927

**Authors:** Hanna Orlikowska-Rzeznik, Emilia Krok, Madhurima Chattopadhyay, Agnieszka Lester, Lukasz Piatkowski

## Abstract

Studies of biological membrane heterogeneity particularly benefit from the use of the environment-sensitive fluorescent probe Laurdan, for which shifts in the emission, produced by any stimulus (e.g. fluidity variations), are ascribed to alterations in hydration near the fluorophore. Ironically, no direct measure of the influence of membrane hydration level on Laurdan spectra has been available. To address this, we investigated the fluorescence spectrum of Laurdan embedded in solid-supported lipid bilayers as a function of hydration and compared it with the effect of cholesterol – a major membrane fluidity regulator. The effects are illusively similar, hence the results obtained with this probe should be interpreted with caution. The dominant phenomenon governing the changes in the spectrum is the hindrance of the lipid internal dynamics. Furthermore, we unveiled the intriguing mechanism of dehydration-induced redistribution of cholesterol between domains in the phase-separated membrane which reflects yet another regulatory function of cholesterol.

## INTRODUCTION

Water hydrating biological membranes is unequivocally essential for the maintenance of cell viability. Living in peculiar physicochemical cooperation, water stabilizes the structure and dynamics of the lipid bilayer1 and mediates its interactions with other biomolecules,^2^ while lipids affect the spatial arrangement and dynamics of adjacent water molecules.^3^ It is generally accepted that biomembranes exist in a fully hydrated environment, however, it should be noted that cell life also involves the local, temporary membrane dehydration events, such as adsorption of biomacromolecules or lipid bilayer fusion, the latter being a key phenomenon to subcellular compartmentalization, cell growth, neurotransmission, fertilization, viral entry, or exocytosis.^4,5^ Hence, it is clear that the mechanistic understanding of such events requires detailed insights into the local membrane hydration state. Yet, its determination is nontrivial, since the extent to which water interacts with different segments of the lipid bilayer is modulated by various factors such as temperature, the type of lipid headgroup, acyl chain composition, and the phase state of a lipid bilayer.^6,7^ Membrane phase is largely governed by cholesterol (Chol) content, which is a key regulator of acyl chains conformational order and lipid dynamics.^8^ Pure phospholipid bilayers are known to exist either in the solid (gel) or liquid-disordered (L_d_) phase. At sufficient concentration, cholesterol promotes the formation of the intermediate phase known as the liquid-ordered (L_o_) phase, which may coexist with the other two.^9^ The L_o_/L_d_ coexistence, manifested as lateral heterogeneity on a nanometer and micrometer scale, is considered to be the most relevant from the biological perspective.^10,11^ One of the approaches to assess membrane heterogeneity is to employ a fluorescent environmentally sensitive probe immersed in a bilayer, such as the most commonly used Laurdan.^12^ Upon electronic excitation, the Laurdan dipole moment significantly increases, giving rise to the dipolar relaxation of the surrounding molecules. The rearrangement of the immediate environment consumes the energy of the excited Laurdan molecule, manifested as a red shift of the emission spectrum.^13^ This accounts for the extreme sensitivity of Laurdan to the polarity and rate of dipolar relaxation of its immediate environment. In the literature, Laurdan is used to probe the membrane heterogeneity referring, often interchangeably, to different membrane physicochemical properties, including lipid order,^14^ hydration,^15^ or the general term fluidity,^16^ and although these features are related to each other, they are not equivalent. Nevertheless, regardless of the property, any shift in the emission spectrum has been taken as a consequence of alterations in the number and/or mobility of water molecules near the Laurdan’s fluorescent moiety, below the glycerol backbone of the phospholipids. Ironically, despite widespread use for more than four decades, no direct measure of the influence of membrane hydration state on Laurdan spectra has been available.

Here, we investigated, for the first time, the spectral response of Laurdan to dehydration of biomimetic cell membranes, directly compared it with the effect of increasing cholesterol content and elucidated the molecular mechanisms that govern the observed changes. By monitoring the fluorescence spectral characteristics of Laurdan during dehydration of membrane with L_o_/L_d_ coexistence, we unveiled an intriguing mechanism of inter-phase cholesterol redistribution, that is of relevance for membrane-centered cellular events. Our results have important implications for the proper interpretation of data obtained with this and other environmental probes, especially when assessing membrane heterogeneity in living systems, where numerous effects, including local variations in hydration and cholesterol content, can be encountered, often simultaneously.

## RESULTS AND DISCUSSION

Changes in the steady-state fluorescence spectrum of Laurdan embedded in the solid-supported lipid bilayers (SLBs) composed of a pure phospholipid (di14:1-Δ9*cis*-PC) resulting from membrane dehydration are depicted in Figure 1a. Membrane hydration state was varied by applying a drying process with a slow and sequential reduction in the sample environment’s relative humidity (RH).^1^

**Figure 1.**
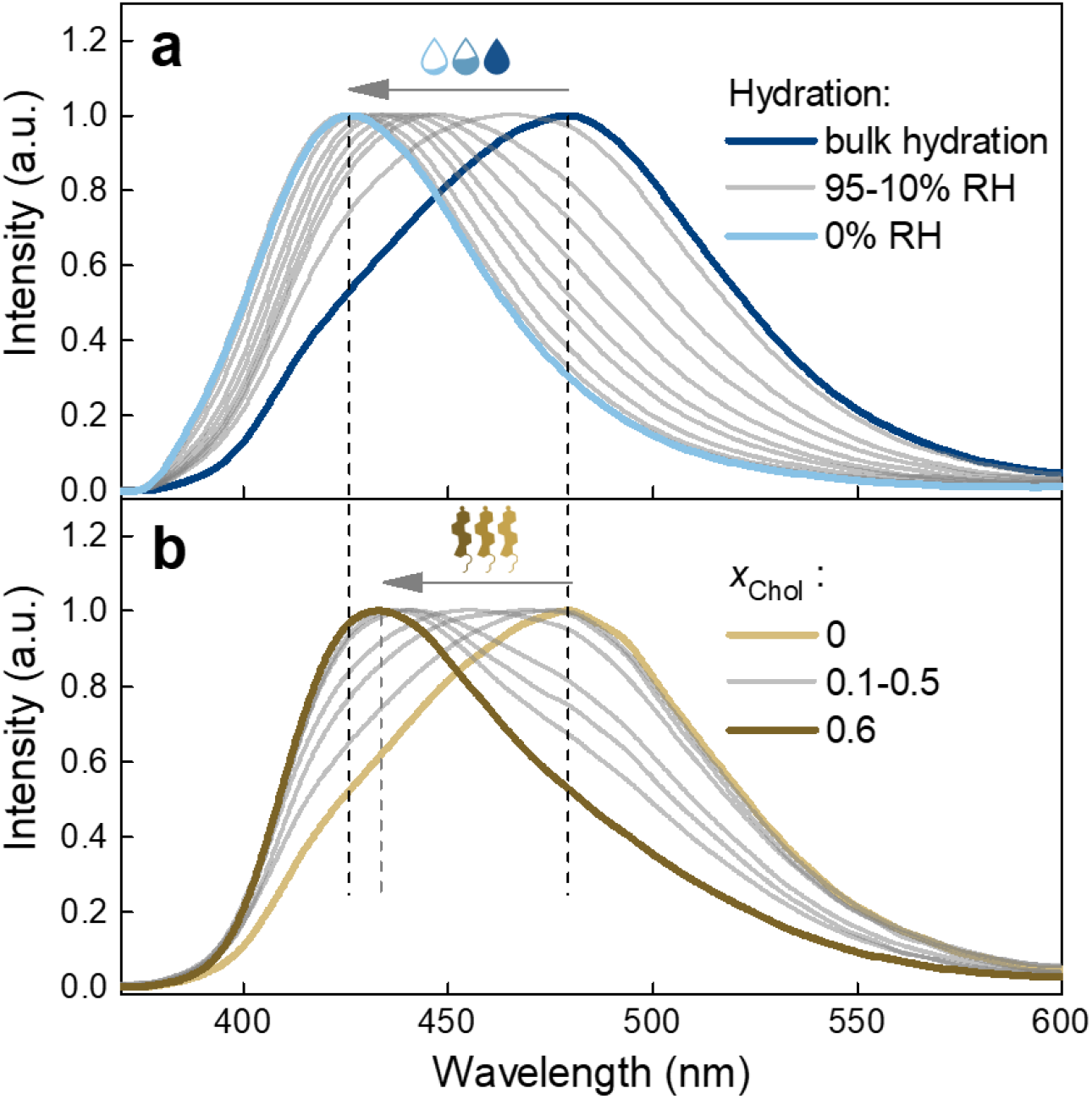
The changes in the fluorescence spectrum of Laurdan embedded in the di14:1-Δ9*cis*-PC SLB induced by; (a) decreasing hydration state from bulk hydration down to 0% RH of the atmosphere surrounding the membrane (the intermediate hydrations are 95, 80, 70, 60, 50, 40, 30, 20 and 10% RH), (b) increasing cholesterol molar fraction *X*_Chol_ from 0 up to 0.6 (the intermediate molar fractions are 0.1, 0.2, 0.25, 0.3, 0.4, 0.5). Laurdan emission spectrum for each hydration step and each *X*_Chol_ is averaged over two different samples, smoothed using fast Fourier transform filter, and normalized.

The fluorescence spectrum of Laurdan in the fully hydrated SLBs is characterized by a broad band with its maximum centered at ~480 nm, a value that is typically attributed to the L_d_ phase,^17^ congruent with the report that at room temperature di14:1-Δ9*cis*-PC lipids form disordered phase.^18^ As the hydration decreases, the spectrum exhibits a progressive blue shift. After drastic dehydration (0% RH), the probe’s emission spectrum resembles that characteristic of ordered membranes (in the gel phase) with the maximum centered at ~430 nm.^19^ Consequently, the observed changes are reflected in the Laurdan generalized polarization (GP), which is a commonly used parameter to assess the overall membrane order (Fig. 2a, blue part).^19^ It is defined as GP = (*I*_440_ - *I*_490_)/(*I*_440_ + *I*_490_), where *I*_440_ and *I*_490_ are the fluorescence emission intensities at 440 and 490 nm, respectively.^20^

**Figure 2.**
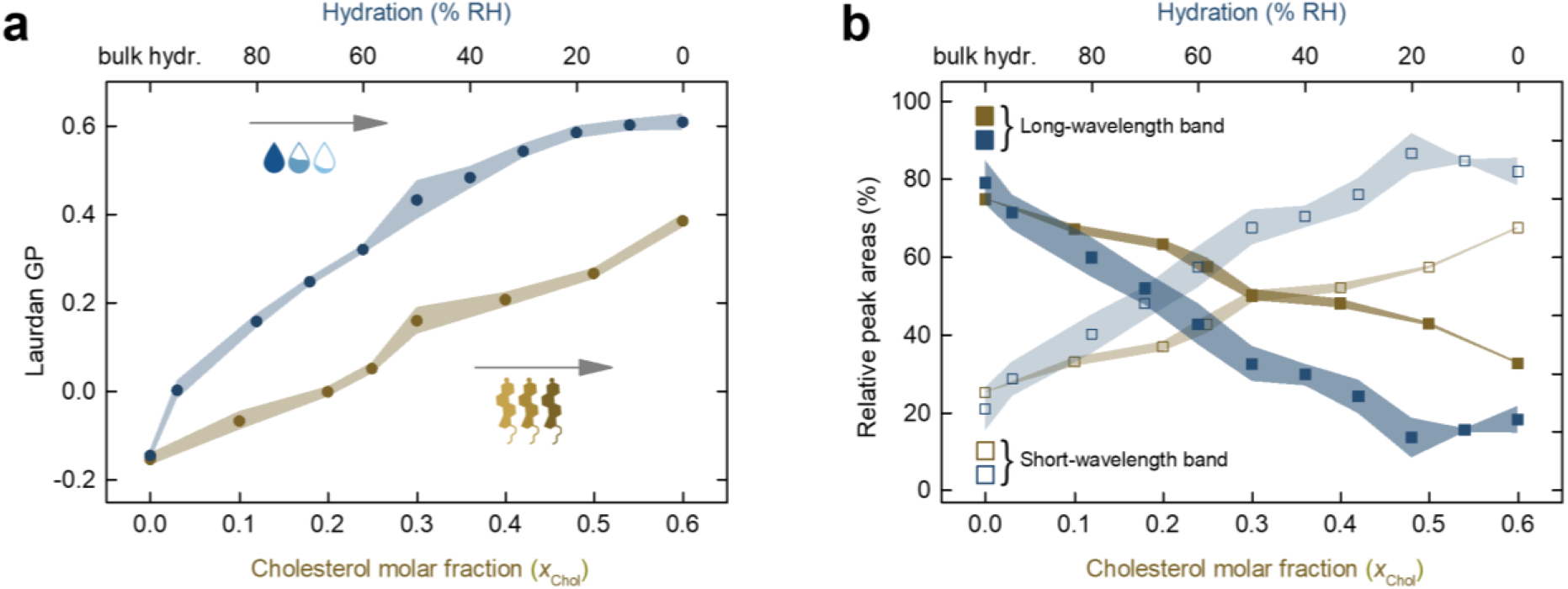
(a) Laurdan GP as a function of membrane hydration and cholesterol molar fraction in a di14:1-Δ9*cis*-PC SLB. (b) The relative area of the two log-normal components that give the best fit to the Laurdan emission spectra in the same membrane system. The uncertainties are standard deviations, denoted as shadows around mean values.

Theoretically, GP can assume values between 1 and −1, however, in the lipid membranes, it does not reach its extreme values and typically scales from 0.6 (the most ordered) to below −0.1 (the least ordered).^21^ As can be seen from Figure 2a (blue part), for a fully hydrated membrane, GP has a negative value, indicative of a disordered bilayer. Upon removal of bulk water and equilibrating the membrane with an atmosphere of 95% RH, the GP takes on a value close to zero and gradually increases with a further reduction of water content, reaching a maximum of 0.62 ± 0.04 for a water-depleted membrane (0% RH), a characteristic of a gel phase.

To confront the pure effect of membrane dehydration with the influence of cholesterol on the Laurdan response, we examined the membranes composed of a binary mixture of di14:1-Δ9*cis*-PC with varying molar fraction of cholesterol (*x*_Chol_). Changes in the probe’s emission spectrum resulting from the increasing *x*_Chol_ are depicted in Figure 1b. The emission peak is significantly affected by the sterol concentration, exhibiting the shift in the same direction as in the case of decreasing hydration. At first glance, changes due to membrane dehydration and due to addition of cholesterol appear very similar. Nevertheless, a few significant differences can be pointed out. Up to *x*_Chol_ = 0.2, the long-wavelength shoulder of the spectrum (~490-550 nm) virtually does not change. Only at *x*_Chol_ = 0.25 and above is a progressive decrease in its intensity observed. In contrast, the lower wavelength part of the spectrum initially exhibits drastic shift and then stops changing above *x*_Chol_ = 0.3. These were not observed in the case of membrane dehydration, for which gradual change of the spectral position was observed throughout the entire dehydration trajectory. Another characteristic feature, evident when analyzing the change in the spectral shape with increasing *x*_Chol_, is a spectral shoulder (~480 nm) that stands out over a wide range of cholesterol concentrations. Most importantly, comparing the extent of the blue shift, it is apparent that membrane dehydration down to 0% RH causes greater changes than the highest cholesterol content *x*_Chol_ = 0.6. The same conclusions can be derived by analyzing the course of the GP parameter (Fig. 2a, brownish part). The increase in cholesterol is accompanied by an increase in Laurdan GP, but to a much lesser extent compared to the pure effect of water depletion, as it reaches a maximum of 0.38 ± 0.03 for the highest *X*_Chol_. We also verified higher molar fractions of cholesterol (0.7 and 0.8), but as expected, they did not produce further changes in the fluorescence spectrum, which is reasonable given the limited solubility of cholesterol in the phospholipid membrane.^22^

It can be noted that Laurdan fluorescence spectrum has a complex line shape, indicative of a heterogeneous local environment. Although due to the dynamic nature of the phospholipid bilayer with existing packing defects,^23^ and the nonuniform, to some extent, insertion depth^24^ and orientation^25^ of Laurdan in the membrane, there may be different subpopulations of Laurdan molecules experiencing distinct environments, we consider that the ability of the environment to adapt (dipolar relaxation) to the excited Laurdan molecule is the major determinant of the Laurdan’s fluorescence spectral properties. Therefore, the Laurdan emission can be modelled in terms of a simple two-state model, assuming that the steady-state spectrum contains two contributions, a short-wavelength band reflecting Laurdan population experiencing little or no dipolar relaxation and a long-wavelength band associated with Laurdan within readily relaxing environment. Thus, to gain further insight into the probe’s local molecular environment upon membrane dehydration and addition of cholesterol, we performed spectra decomposition using two log-normal line shapes as proposed by Bacalum et al.^26^ (Fig. S1 and Materials and Methods; fluorescence spectra analysis section in SI). The results of such an analysis are plotted in Figure 2b as the relative areas of the short-wavelength and long-wavelength bands, reflecting the percentages of Laurdan populations associated with non-relaxed and relaxed local solvent environment, respectively, as a function of the membrane hydration (blue part) and cholesterol content (brownish part). The exemplary extracted spectra for different cholesterol molar fractions are shown in Figure S2.

The Laurdan fluorescence spectra, at all degrees of hydration, can be well described by a superposition of two log-normal line shapes, with peaks around 475 and 427 nm, confirming that the fluorescence decay is mainly due to transitions from only two different excited energy levels. In a pure phospholipid bilayer under fully hydrated conditions, the most (~79%) of the fluorescence emission is due to the long-wavelength transition of Laurdan residing in hydrated, relaxed environment (Fig. 2b, blue part). As water content is lowered, the population of Laurdan molecules that experiences a dipolar relaxation starts decreasing, giving rise to the short-wavelength transition. A steady decrease of the population within relaxed environment (and concomitant increase of population experiencing non-relaxed medium) is observed from a fully hydrated state to around 20% RH. Below this value, the band fractions reach a plateau with the relative populations of the two Laurdan populations accounting for ~18% and ~82%, respectively, so the opposite of full hydration.

Noticeably, cholesterol produces more subtle changes (Fig. 2b, brownish part). Analogously to the dehydration process, the Laurdan fluorescence spectra, at all *X*_Chol_, can be well reconstructed by a superposition of two log-normal lines, with maxima around 482 and 430 nm (Figure S2). Throughout the analyzed range of cholesterol concentrations, the differences in the proportion of the populations emitting from within the relaxed and non-relaxed solvent environments are not as pronounced compared to the effect of decreasing membrane hydration. In other words, compared to dehydration, even at high *X*_Chol_, the considerable number of Laurdan molecules experience a dipolar relaxation.

Now, let us consider the physical origin of the observed changes. The first thing that comes to mind, when observing the shift of the spectrum toward shorter wavelengths as the membrane is dehydrated, is the gradual reduction in the amount of water molecules around the fluorophore. It must be emphasized, however, that Laurdan fluorescent moiety localizes below the glycerol backbone of phospholipids, near the *sn-1* carbonyls,^27^ where water molecules are scarce and strongly bound via hydrogen bonds to the lipid carbonyl oxygen atoms.^28–30^ Therefore, we assume the Laurdan emission to be sensitive to the changes in its direct vicinity, rather than at the level of phosphates or even the more outer parts of the membrane. In the lipid membrane interphase region, beyond the carbonyls, water is distributed around the phosphate and choline groups. It has been reported that, on average, carbonyl oxygen atoms form 1 H-bond per lipid, phosphate non-estrified oxygen atoms form 4 H-bonds and other lipid oxygen atoms form 1 H-bond.^28^ Yet, the choline moiety, due to the non-polar character of the methylenes, cannot form H-bonds with adjacent water molecules. Instead, it organizes the water molecules via weak electrostatic and van der Waals interactions so that they form a clathrate structure around it, containing, in case of a zwitterionic phosphocholine lipid, about 6 water molecules.^31^ In total, these 12 water molecules are considered a first hydration shell. Subsequent hydration shells incorporate exclusively water molecules that are unbound to lipids and are assumed to be localized mostly in the outer parts of the membrane.^32^ During dehydration, it is the strength of intermolecular interactions that governs the order of desorption of water molecules. As such, loose water molecules interacting only with each other, through relatively weak hydrogen bonds, along with the water molecules directly associated with phospholipids via the weak van der Waals interactions are removed first. These are followed by desorption of water molecules bound more strongly to polar residues of phospholipids.^32^ It has been shown that upon removal of bulk water and exposing the lipid bilayers to 95% RH, the first solvation shell is largely preserved and only further reduction of hydration degree breaks it down.^1^ However, even extreme dehydration does not remove water strongly bound to lipids, particularly these associated with the carbonyls.^31^ Having this molecular picture in mind, we infer that upon membrane dehydration, the most drastic changes in hydration occur in the outer regions of the phospholipid bilayer, while the number of water molecules in the vicinity of Laurdan fluorophore barely changes. Thus, the rationale behind Laurdan response must be different. Importantly, the nanosecond solvent relaxation kinetics revealed by the time-dependent fluorescence shift measurements of Laurdan in phospholipid bilayers, is associated with the collective rearrangement of the hydrated *sn-1* carbonyls and not water molecules itself.^33^ In addition, it was demonstrated that GP calculated from the steady-state Laurdan emission spectra correlates well with rearrangement kinetics of the immediate vicinity of the fluorophore and not the total spectral shift (which mirrors the polarity and thus the number of water molecules).^33^ This implies that Laurdan GP primarily reflects the mobility of hydrated functional groups of lipids at the Laurdan level rather than the extent of water penetration. The mobility of lipid carbonyls, in turn, has been found to be dependent on the local hydrogen bond network dynamics.^34^ Noteworthy, both experimental and molecular dynamics (MD) simulation studies unveiled the slowdown of interfacial water dynamics induced by membrane dehydration.^29,35^ The results indicate an increasing residence time of bound water molecules within the lipid polar groups as water content decreases. The more persistent the hydrogen bonds between water molecules and carbonyl oxygens, the more restricted dynamics of hydrated carbonyls. Furthermore, in addition to the slowdown of interfacial water dynamics, both the structural and dynamical properties of lipid bilayers have been found to be affected by water content.^1,35–37^ Membrane dehydration results in a decrease in the area and volume per lipid and a concomitant increase in membrane thickness, as well as a slowdown in the lipid translational and rotational mobility, ultimately leading to a liquid-disordered to gel phase transition. The results of the log-normal decomposition reinforce the idea that during membrane dehydration there is no significant change in the hydration level at the Laurdan site. Had the number of water molecules aligning around the Laurdan dipole decreased, indicating a decrease in the polarity of the immediate vicinity of the fluorophore, a shift in the peak wavelength would have been observed. Instead, we obtained stable positions of the peaks (see figure S1). Taken together, we interpret the decrease in the area of the long-wavelength band as a diminishing population of Laurdan molecules, for which the collective relaxation of the hydrated lipid groups completes within Laurdan’s fluorescence lifetime. In other words, as the water depletion in the bilayer progresses, the number of localized sites where hydrogen bond network and lipid dynamics allow for the hydrated carbonyls to reorient along the Laurdan excited state dipole decreases.

The influence of membrane dehydration and the effect of cholesterol on the Laurdan emission spectrum are illusively similar, but not equivalent. Importantly, congruent with our spectral decomposition results, as suggested by Amaro et al.,^38^ the presence of cholesterol does not affect the polarity (number of water molecules) in the vicinity of Laurdan fluorophore. Therefore, cholesterol-induced changes in the probes’ emission spectrum must be due to the reduced kinetics of dipolar relaxation. Both experimental and theoretical investigations revealed that contrary to dehydration, an increase in cholesterol content in the membrane accelerates the relaxation of interfacial hydrogen-bond network.^39–41^ According to the theoretical predictions, the incorporation of cholesterol into the phospholipid bilayer leads to a rupturing of inter-lipid H-bonds bridging two adjacent phospholipids, which are considered more rigid and slower, and an accompanying increase in the fraction of lipid-water H-bonds, which are faster and more mobile. Collectively, it leads to an enhancement in the water mobility at the interface. This effect rather promotes the dipolar relaxation around the Laurdan fluorescent moiety, therefore, there must be different dominant effect that leads to the overall blue shift. Cholesterol is known to induce phospholipid bilayer ordering, as manifested by a significant increase in the C–H bond order parameter of different segments in the acyl chains of lipids in nuclear magnetic resonance experiments.^42–44^ However, it should be emphasized that this relates only to the hydrophobic core of the membrane. The structural order parameters of the interfacial regions of the phospholipid bilayer, namely the choline, phosphate, and glycerol backbone of the lipid headgroups along with the carbonyl region, remain virtually unaffected by the presence of cholesterol.^43,44^ Therefore, it is highly unlikely that it is the structural conformational ordering that causes such drastic changes in the Laurdan spectrum upon addition of cholesterol. It is worth noting, however, that the conformational order reflects the orientation of the C–H bond vector with respect to the bilayer normal averaged over the lipid ensemble and over time,^45^ but does not carry information about its dynamics. Recent work by Antila et al.^44^ unveiled that although cholesterol causes only marginal changes in the structural order of the membrane region where Laurdan resides, it significantly impedes the dynamics of the glycerol backbone and the associated carbonyls. This implies that the predominant phenomenon governing the cholesterol-induced blue shift of the Laurdan fluorescence spectrum is the slowing down of intralipid dynamics.

After determining both the effect of dehydration of a pure phospholipid bilayer and cholesterol incorporation, we evaluated the Laurdan response to dehydration of a phase-separated membrane, which is considered a much better mimic of biological membranes. To this end, we used an equimolar ternary mixture of di14:1-Δ9*cis*-PC, cholesterol and egg sphingomyelin (eggSM), which at the room temperature exhibits the L_o_/L_d_ phase coexistence (Fig. S3). The use of SLBs as samples and fluorescence microscopy coupled to a spectral detection enabled collection of spectra separately from the L_d_ and L_o_ domains. Under fully hydrated conditions, emission of Laurdan in L_o_ is blue-shifted and significantly narrower than for Laurdan in the L_d_ phase (Fig. S4), consistent with the previous observations.^46^ As expected, as the hydration level decreases the fluorescence spectrum of Laurdan in L_d_ shifts toward shorter wavelengths (Fig. S4a). Changes for L_o_ phase are much less pronounced (Fig. S4b and S5), therefore we focus here on the L_d_. The discussion on the insensitivity of Laurdan to dehydration of the L_o_ phase can be found in the supplementary info (SI Note 1). Complete dehydration of L_d_ domains resulted in a smaller shift in the spectrum than dehydration of the single-component membrane, but greater than for a membrane with the maximum cholesterol content. It is worth noting that the lateral organization of the membrane was monitored between the spectra collection routes for distinct hydration states and it was confirmed that the phase separation of the membrane remained virtually unaltered during the dehydration process (see Fig. S3). Comprehensive data on this issue can be found in our previous work.^1^

Analysis of the GP parameter as a function of hydration level of L_d_ domains reveals interesting behavior (Fig 3a, green part).

**Figure 3.**
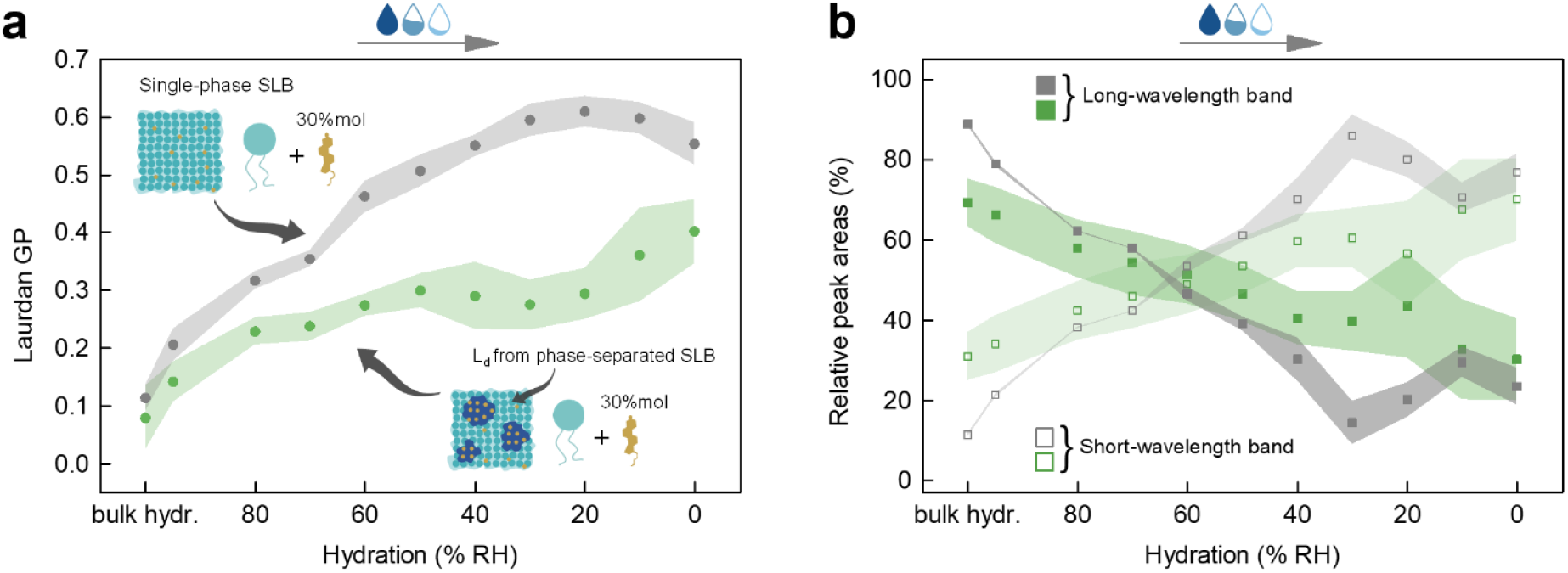
(a) Laurdan GP for L_d_ domains from phase-separated SLB composed of an equimolar mixture of di14:1-Δ9*cis*-PC/Chol/eggSM and for the counterpart from SLB without phase separation composed of a binary mixture of di14:1-Δ9*cis*-PC/Chol with *X*_Chol_ = 0.3 as a function of hydration level. (b) The relative area of the two log-normal components that give the best fit of the Laurdan emission spectra in the same membrane systems as a function of hydration. The uncertainties are standard deviations, denoted as shadows around mean values.

In the range from bulk hydration to 50% RH, a gradual, small increase in GP can be observed, after which it remains constant down to 20% RH, and then it increases slightly again. It is qualitatively different than in the case of pure phospholipid SLBs dehydration. Intrigued by this, we examined whether changes in GP caused by the dehydration process of a membrane without phase separation, but with the same composition as in the L_d_ domains, occur in the same way. By comparing the line shape of the Laurdan fluorescence spectra acquired from the L_d_ domains with the ones originating from the monophasic membranes with different *X*_Chol_, we inferred that under fully hydrated conditions, the *X*_Chol_ in L_d_ is in the range of ~0.25-0.3. Therefore, to reproduce the molecular L_d_ composition in the membrane without phase separation, we prepared monophasic lipid membranes with *X*_Chol_ = 0.3 and yet again monitored the Laurdan fluorescence during the dehydration process. It was assumed that since the two systems at bulk hydration are compositionally the same, the dehydration process would cause the same changes in Laurdan emission. Changes of the Laurdan spectrum in a membrane composed of a binary mixture of di14:1-Δ9*cis*-PC and Chol due to dehydration are demonstrated in Figure S6. The course of the GP value as a function of hydration state of this membrane (Fig. 3a, gray part) qualitatively resembles that for a pure phospholipid membrane (Fig. 2a), except that it starts from a slightly higher value at full hydration (indicative of increased order) and stops changing below 30% RH, but reaches the same average value of 0.62. But most importantly, and surprisingly, it does not resemble the trajectory for its counterpart from the phase-separated membrane (Fig. 3a, green part). For membrane without L_o_ domains present, the changes are steeper and do not exhibit a plateau in the range from 50 to 20% RH. In addition, in general, GP has significantly higher values for each of the hydration levels, except for bulk hydration, compared to the L_d_ domains from phase-separated membrane. The spectral global analysis further highlights these differences (Fig. 3b). The results are rather intriguing and indicative of an additional mechanism that counteracts and effectively softens the changes caused by dehydration of L_d_ in phase-separated membrane. The observed trajectories (whether GP or resulting from spectral decomposition) resemble those observed for changing cholesterol content (Fig. 2b) rather than those for dehydration, suggesting that perhaps cholesterol content in L_d_ phase changes. This is feasible as L_o_ phase contains more cholesterol (~70% of all cholesterol in the membrane) and may act as a reservoir of cholesterol in the phase-separated membranes. Therefore, we reason that the peculiar dehydration-induced behavior of L_d_ phase in phase-separated membrane might be due to redistribution of components between L_o_ and L_d_ domains, most likely involving cholesterol. This would rationalize also our previous findings,^1^ that with reduced hydration, the hydrophobic mismatch between the L_d_ and L_o_ domains decreases significantly (cholesterol influx to L_d_ phase would increase its thickness, thus lessening the hydrophobic mismatch between domains). To test our hypothesis, we conducted the fluorescence microscopy experiments with fluorescently labeled cholesterol, the results of which are depicted in Figure S7. Under fully hydrated conditions, as expected, higher intensity of TopFluor-Chol emission is found in L_o_ domains. However, with reduced hydration, the contrast diminishes until it becomes reversed, showing that the L_d_ domains contain more cholesterol. Therefore, it can be concluded that cholesterol influx from L_o_ to L_d_ phase counteracts the dehydration-induced extensive changes in fluidity. Though, the study of cholesterol migration was not the aim of this work and an in-depth understanding and quantification of this phenomenon require additional experiments and analysis.

## CONCLUSIONS

In conclusion, we have shown that the effects of membrane dehydration and cholesterol incorporation on Laurdan’s fluorescence spectrum are illusively similar, thus interpretation of data obtained with this probe should be done with caution. We evidence that the dehydration-induced changes in the Laurdan’s emission spectrum result from the conformational ordering of lipids and hindrance of the lipid internal motions along with the slowdown of hydrogen bond network dynamics acting collectively to impede the dipolar relaxation around the probe’s excited state dipole. In the case of cholesterol incorporation, for which neither hydrogen bond network relaxation slowdown nor static conformational ordering of the lipid bilayer region probed by Laurdan is observed, changes in the emission are likely caused only by the hampered dynamics of the glycerol backbone and the associated carbonyls, which rationalizes the more subtle changes compared to membrane dehydration. Furthermore, we have discovered that the dehydration of the phase-separated membrane drives the redistribution of cholesterol between domains, which likely acts as a regulatory mechanism to prevent excessive deviations in fluidity that may destabilize the cell membrane and hence be harmful to the cell. This intriguing finding adds to the multiple actions of cholesterol towards mechanochemical homeostasis of the lipid membranes. Our results provide new insights at the intersection of physical chemistry, photo- and biophysics and should stimulate the design of a range of new experiments and simulations regarding the specificity and sensitivity of environmental probes.

### Supporting Information

Materials and Methods, Supplementary Experimental Results; Additional results of the independent and global fitting procedure for different SLB systems, Fluorescence microscopy images of a phase-separated SLB as a function of membrane hydration state, Fluorescence spectra of Laurdan in liquid-disordered and liquid-ordered domains from phase-separated SLB and in one-phase SLB with molar fraction of cholesterol 0.3 as a function of membrane hydration state, Generalized polarization of Laurdan in liquid-ordered phase as a function of membrane hydration (PDF).

## Notes

The authors declare no competing financial interests.

## ACKNOWLEDGMENTS

This work was financed from the budget funds allocated for science in the years 2019– 2023 as a research project under the “Diamond Grant” program (decision: 0042/DIA/2019/48). The authors also acknowledge the financial support from the EMBO Installation Grant 2019 (IG 4147) and National Science Centre (Poland) 2020/37/B/ST4/01785. L.P. acknowledges the financial support from the First TEAM Grant No. POIR.04.04.00-00-5D32/18-00, provided by the Foundation for Polish Science (FNP).

