## Supplementary Information for "Laurdan discerns lipid membrane hydration and cholesterol content"

### Materials and Methods

#### *Materials*

Lipids 1,2-dimyristoleoyl-sn-glycero-3-phosphocholine (di14:1-Δ9*cis*-PC), egg yolk sphingomyelin (eggSM), cholesterol (Chol), and 23-(dipyrrometheneborondifluoride)-24-norcholesterol (TopFluor-Chol), were supplied by Avanti Polar Lipids (Alabaster, AL, USA). Fluorescent probe 6-dodecanoyl-2-dimethylaminonaphthalene (Laurdan), phospholipid 1,2-dioleoyl-sn-glycero-3-phosphoethanolamine labeled with Atto 633 (Atto 633-DOPE), monosialoganglioside (GM1) from bovine brain, and chloroform (HPLC grade) were purchased from Merck KGaA (Darmstadt, Germany). Alexa Fluor 594 conjugated with cholera toxin subunit B (Alexa Fluor 594-CTxB) were obtained from Molecular Probes, Life Technologies (Grand Island, NY, USA). Buffer reagent 4-(2-hydroxyethyl)piperazine-1-ethanesulphonic acid (HEPES PUFFERAN®) was obtained from Carl Roth GmbH + Co. KG (Karlsruhe, Germany). Calcium chloride ($CaCl_2$) was purchased from Chempur (Piekary Slaskie, Poland). Sodium chloride (NaCl) was supplied by PPH STANLAB Sp. z o.o. (Lublin, Poland). All compounds were used without further purification. Ultrapure water was acquired using the Milli-Q Direct Water Purification System from Merck KGaA (Darmstadt, Germany). Optical adhesive UV-activated glue Norland 68 was purchased from Thorlabs Sweden AB (Mölndal, Sweden). The sheets of mica used for the preparation of solid supports for the lipid membranes were obtained from Shree GR Exports Private Limited (Kolkata, India). Glass coverslips No. 0 were purchased from Paul Marienfeld GmbH & Co. KG (Lauda-Königshofen, Germany).

#### *Solid-supported lipid bilayers fabrication*

Solid-supported lipid bilayers (SLBs) were prepared using vesicle deposition on a solid substrate procedure as described previously[1] with appropriate modifications. SLBs of different compositions were examined: (i) pure di14:1-Δ9*cis*-PC, (ii) binary mixtures di14:1-Δ9*cis*-PC/Chol with varying Chol molar ratio ($x_{Chol}$ = 0.1; 0.2; 0.25; 0.3; 0.4; 0.5; 0.6), and (iii) an equimolar ternary mixture di14:1-Δ9*cis*-PC/Chol/eggSM. First, the membrane components were mixed along with the fluorescent probe(s) to form a chloroform solution of the specified composition with a final lipid concentration of 10 mM. The lipid to each fluorescent probe molar ratio was 1000:1. For the fluorescence spectra measurements, two probes – Laurdan and Atto 633-DOPE were used, whereas, for the confocal microscopy experiments three probes – Atto 633- DOPE, TopFluor-Chol, and Alexa Fluor 594-CTxB-GM1 complex were used to label the membrane. The appropriate solution was then dried under nitrogen gas, followed by desiccation in the vacuum chamber for at least 2 hours. The lipid film was then hydrated in buffer solution (10 mM HEPES and 150 mM NaCl, pH adjusted to 7.4) to obtain a 10 mM lipid concentration. The lipid suspension was subjected to four cycles of heating to 60°C and vortexing, with each heating and vortexing step taking 1 minute, producing multilamellar vesicles (MLVs). The lipid mixture was diluted 10-fold in a buffer to yield a 1 mM MLV suspension, and then distributed into sterilized glass vials and stored at -20°C for further use. The aliquoted MLV suspension of the desired composition was bath-sonicated for at least 10 minutes until the solution became transparent, indicating the formation of small unilamellar vesicles (SUVs). To prepare a solid support for SUVs deposition, a small amount of immersion oil was deposited onto glass coverslip No. 0, over which a thin sheet of freshly cleaved mica, cut beforehand as round plates with a diameter of 9.53 mm (3/8 inches), was placed and adhered with UV-activated glue around the periphery of the substrate. A microcentrifuge tube's lid and bottom were cut off and the resulting cylinder was placed on a coverslip and sealed with silicone to form a reservoir with mica at the bottom. 100 µl of SUVs suspension was deposited on the mica surface followed by the immediate addition of 2 µL of 0.1 M $CaCl_2$ solution. After 30 seconds, 600 µL of the previously used buffer solution was added to prevent the hydration layer from drying out. After 30 minutes of incubation at ambient temperature, the SLB was rinsed 10 times with 2 mL of buffer solution to wash out excess, unburst vesicles. Finally, the remaining volume of the tube was filled with buffer solution, and this condition is called the fully hydrated state of the membrane throughout the paper.

#### *SLB hydration state control*

To perform a direct measurement of the effect of the hydration state of the lipid bilayer on the Laurdan fluorescence spectrum, we employed our home-built humidity control set-up,[1,2] assuring a controlled drying process with a slow and sequential decrease in relative humidity (RH) of the membrane environment. The set-up consists of a nitrogen gas ($N_2$) cylinder, 3 flow meters, 3 manual valves, a reservoir with water (for water-vapor-saturation), and an electronic hygrometer. In brief, to reduce the SLB hydration, bulk water was first removed with a micropipette from the sample container until no buffer droplets on the mica surface were visible to the naked eye. Nitrogen gas of 95% RH was then immediately, gently blown into the sample container. The relative humidity of $N_2$ was regulated by mixing of wet (water-vapor-saturated, 95% RH) and dry (0% RH) gas streams. Wet and dry $N_2$ gas flows were individually regulated by two manual valves, while monitoring the readings of two flow meters connected to the flow paths. A third flowmeter and manual valve were used to keep the final $N_2$ gas flow rate constant at ~1.2 L/min throughout the experiment. The electronic hygrometer allowed monitoring of the relative humidity and temperature of the final gas flow, indicating the possible need for adjustment. The dehydration was performed from 95 to 80% RH and further in steps of ~10% RH to 0% RH. The SLB atmosphere was equilibrated to the specified relative humidity after about 10 minutes and only then were the sample imaged and Laurdan emission spectra recorded.

#### *SLB imaging and steady-state emission spectra acquisition*

The main experiments were carried out on a manual, inverted microscope (Carl Zeiss, Axiovert 200). The excitation beam at 370 nm was provided by a pulsed supercontinuum laser (NKT Photonics, SuperK FIANIUM FIU-15) equipped with a UV extension unit (NKT Photonics, SuperK EXTEND-UV). In all of the experiments described, we used nonpolarized excitation. A 50/50 beam splitter was used to reflect the excitation light into an oil immersion objective (Carl Zeiss, EC Plan-Neofluar 40x/1.30), which focused the beam to a diffraction-limited spot in the sample plane. Epifluorescence signal was spectrally filtered using a 380 nm long-pass filter (Semrock, FF01-380/LP-25) and guided to a single photon counting module (Hamamatsu Photonics, C11202-100) for imaging purposes or to a spectrograph (Andor, Kymera 328I-C), where it was spectrally dispersed with a 150 lines/mm grating and subsequently detected with an electron multiplying charge-coupled device camera (Andor, iXon 888 UCS-BB), pre-cooled to −70 °C, for spectral measurements. One or the other detection path was selected with the help of a remotely controlled mirror. Single photon counting module counts were read and converted to a digital signal by data acquisition card (National Instruments Corporation, NI USB-6363). The sample was scanned across the fixed laser foci with a piezoelectric nanopositioning stage (Mad City Labs, Nano-LPS200) in x-y dimension. Nano-Drive 3 controller (Mad City Labs) was used for controlling the scanning stage. Image reconstruction and positioning of the sample were controlled using a home-made LabVIEW program. To avoid excessive photobleaching, sample illumination was synchronized with data acquisition using an optical beam shutter (Thorlabs, SHB1T).

Monitoring of the cholesterol distribution in the SLB with multiple probes as a function of membrane hydration was realized using laser-scanning confocal upright microscope (Carl Zeiss, LSM 710) with an oil immersion objective (Carl Zeiss, EC Plan-Neofluar 40x/1.30). Lasers of wavelengths 633, 488, and 543 nm were used for excitation of Atto 633-DOPE, TopFluor-Chol and Alexa Fluor 594-CTxB-GM1, respectively. The laser power was adjusted during imaging to avoid excessive photobleaching of the sample.

#### *Fluorescence spectra analysis*

Laurdan generalized polarization (GP) was calculated from the equation introduced by Parasassi et al.[3] and most commonly used in the literature: $GP = \frac{I_{440} - I_{490}}{I_{440} + I_{490}}$, where $I_{440}$ and $I_{490}$ are fluorescence intensities averaged over 5 data points (~2 nm) around 440 nm and 490 nm, respectively. The averaging was done to compensate for the noise present in the spectra. Each GP value demonstrated in the figures is averaged over at least 10 different spots from each of the samples at a particular membrane hydration state or each cholesterol molar fraction (the number of samples varies from experiment to experiment and ranges from 1 to 4). The uncertainties were calculated as standard deviations.

Spectral decomposition of the fluorescence spectra acquired for the samples with specific composition and at specific conditions was done using two log-normal functions in the form[4]:

$$\begin{cases} I = I_m exp\left[-\frac{ln2}{ln^2(\rho)} ln^2\left(\frac{a-\upsilon}{a-\upsilon_m}\right)\right] & if\ \upsilon < a \\ I = 0 & if\ \upsilon > a \end{cases} \tag{1}$$

where $I$ represents the fluorescence emission intensity, $I_m$ is the maximum of intensity, $\upsilon$ is the wavenumber, $\upsilon_m$ is the spectral position of the maximum intensity of the log-normal function, $\rho = \frac{\upsilon_m - \upsilon_{min}}{\upsilon_{max} - \upsilon_m}$ is the asymmetry of the function ($\upsilon_{max}$ and $\upsilon_{min}$ represent the wavenumber values at half intensity), and $a$ is the limiting wavenumber: $a = \upsilon_m + \frac{(\upsilon_{max} - \upsilon_{min})\rho}{\rho^2 - 1}$. First, fluorescence spectra (averaged from at least 5 spots from each of the samples at a particular membrane hydration state or each cholesterol molar fraction) were fitted to a sum of two log-normal functions, independently for each sample and for each hydration/cholesterol content. In the fitting procedure, performed in Matlab, emission intensity $I_m$, spectral position of the maximum intensity of the log-normal function $\upsilon_m$, as well as spectral positions determining the asymmetry of the function $\upsilon_{max}$ and $\upsilon_{min}$ were all kept as free parameters. The values for all the parameters were restricted to take up physically meaningful values. To ensure that the two log-normal functions do not exhibit excessive asymmetry, we restricted $\upsilon_m - \upsilon_{min}$ to take up values no larger than 1.5 times $\upsilon_{max} - \upsilon_m$. All fitted parameters took values within the imposed bounds for over 90% of the fitted spectra. We point out that throughout the manuscript and supplementary information all spectral data are presented in the wavelength space - experimental data are acquired in the wavelength space, thus such a representation is more intuitive and also can be easily compared with other literature data showing Laurdan fluorescence spectra.

From the individual spectral fits it was evident that the two log-normal functions (referred to as short-wavelength and long-wavelength band in the main text) describe all the acquired spectra very well (see Figure S1a,b), yielding high values of the coefficient of determination ($R^2 > 0.993$ for all fitted spectra). Moreover, we note that spectral position of the maximum intensity ($\upsilon_m$) of each fitted function did not exhibit significant changes as a function of hydration/cholesterol content, clearly pointing at the interconversion of the two (short wavelength/long wavelength) populations (see figure S1c,d). The observed frequency shifts of the two bands for all hydration/cholesterol conditions are small (~5-10 $cm^{-1}$) with respect to the separation between the bands (>50 $cm^{-1}$) and are random rather than showing a specific trend.

Next we performed global fits (n spectra for all hydration conditions or cholesterol content for each sample with the specific composition), in which $\upsilon_m$ was kept as a global parameter for each of the two bands. All other parameters were allowed to vary for each sample condition. We used averaged parameter values from individual fits as starting parameters for the global fit. An exemplary result of the global fit is shown in Figure S2.

The populations of Laurdan experiencing solvent relaxation and of Laurdan embedded in non-relaxing environment were obtained by integrating the short-wavelength and long-wavelength bands, respectively and represented as band areas relative to the area of the entire fluorescence spectrum [%].

### Supplementary Experimental Results

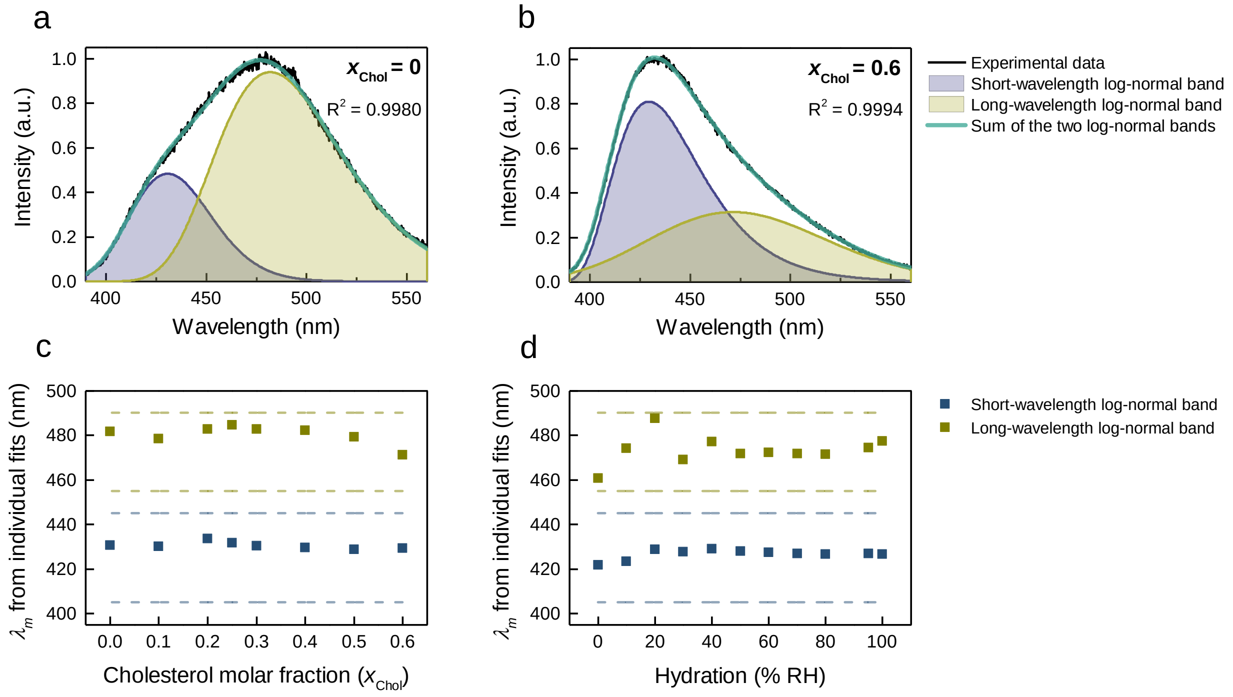


**Figure S1.** The exemplary results of the independent, individual spectra fitting procedure. Two-peak log-normal decomposition of the fluorescence spectra of Laurdan in an exemplary SLB composed of (a) pure di14:1-*Δ9cis*-PC and (b) the binary mixture of di14:1-*Δ9cis*-PC/Chol at $x_{\mathrm{Chol}} = 0.6$ under fully hydrated conditions. Coefficients of determination $R^2$ are indicated in the figures. Spectral positions of the maximum intensity $\lambda_m$ of the two log-normal functions as a function of (c) $x_{\mathrm{Chol}}$ for an exemplary single-phase SLB and (d) membrane hydration of the liquid-disordered phase for an exemplary phase-separated SLB. Blue and olive dashed lines correspond to the limiting range in the fitting procedure for short-wavelength and long-wavelength bands, respectively.

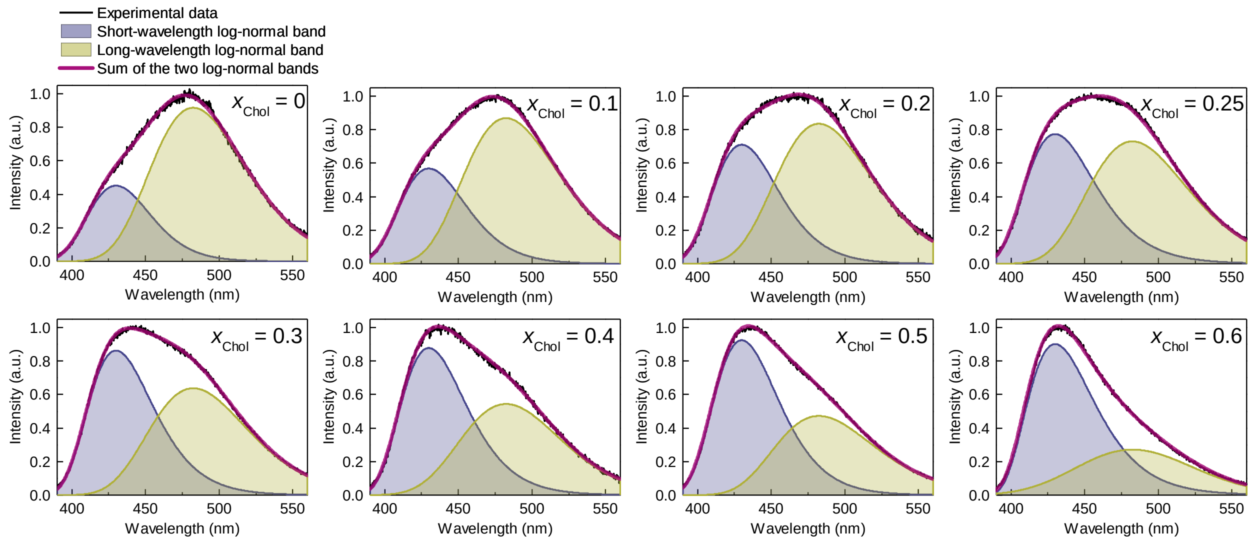


**Figure S2.** An exemplary result of the global fitting procedure. Two-peak log-normal decomposition of the fluorescence spectra of Laurdan in an exemplary SLB composed of di14:1-*Δ9cis*-PC and different cholesterol molar ratio $x_{\mathrm{Chol.}}$ Coefficient of determination $R^2 = 0.9990$.

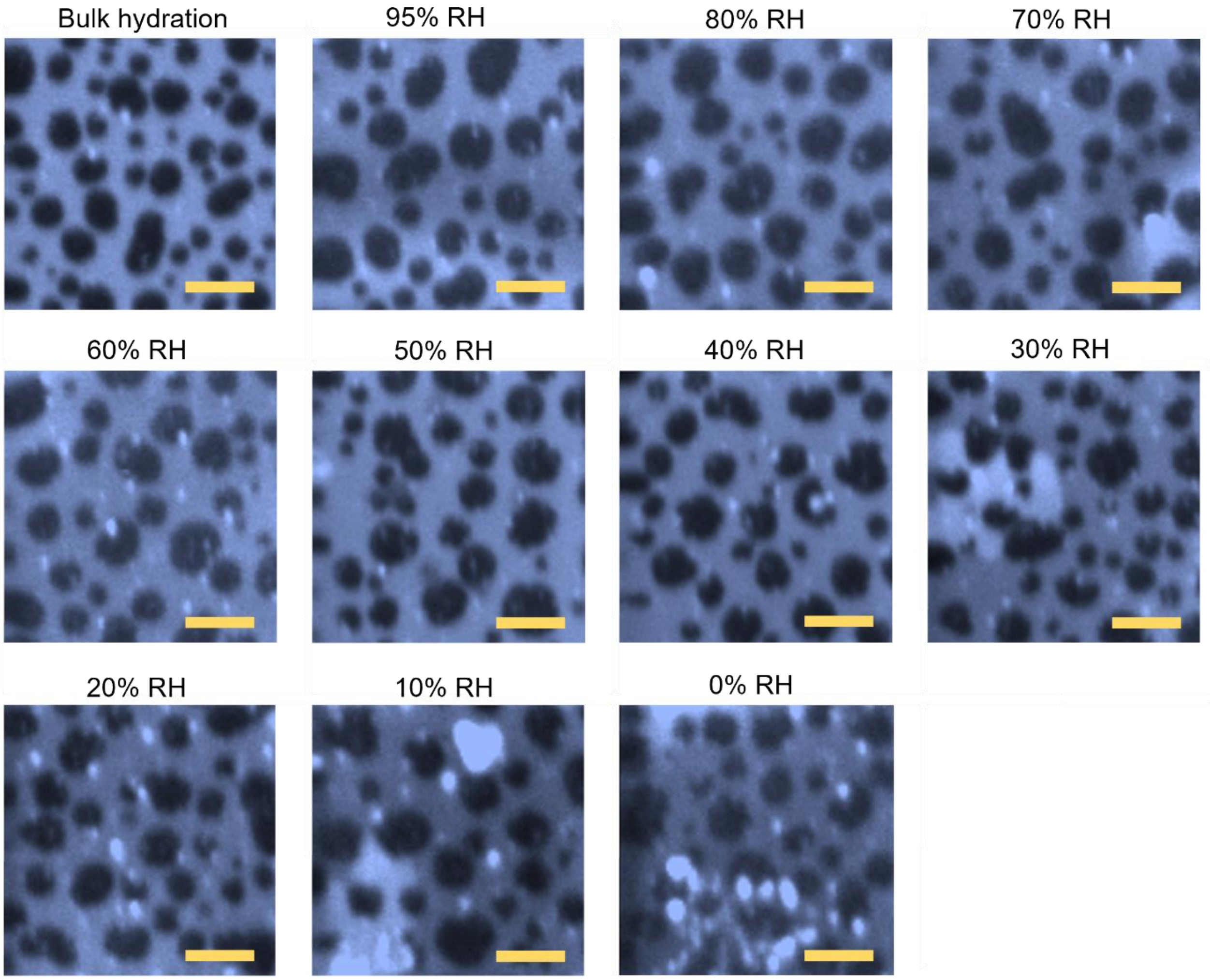


**Figure S3**. Fluorescence microscopy images of an exemplary phase-separated solid-supported lipid bilayer composed of an equimolar mixture of di14:1-Δ9*cis*-PC, cholesterol, and egg sphingomyelin as a function of membrane hydration state. The membrane exhibits phase separation into liquid-disordered (bright regions) and liquid-ordered (dark regions) domains. The membrane was labeled with Laurdan, which distributes evenly in the membrane regardless of phase, and Atto 633-DOPE, which localizes mainly in the liquid-disordered phase, and from this comes the contrast. The concentration of each dye was 0.1% mol. Each image represents a different area within the sample. The images shown originate from one of the four phase-separated samples analyzed in this work. The scale bar corresponds to 5 μm.

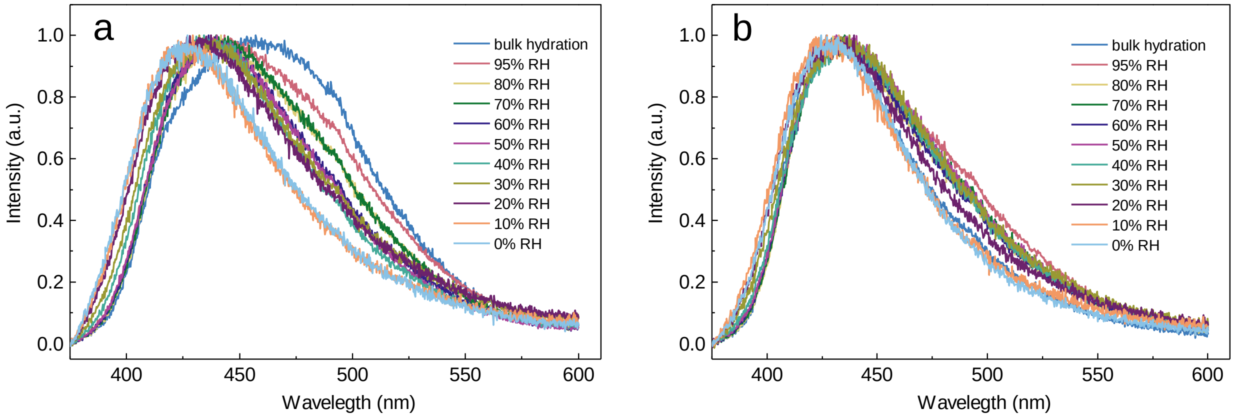


**Figure S4.** Fluorescence spectrum of Laurdan embedded in an exemplary phase-separated solid supported lipid bilayer composed of an equimolar mixture of di14:1-Δ9*cis*-PC, cholesterol, and egg sphingomyelin as a function of membrane hydration state collected from (a) liquid-disordered domains and (b) liquid-ordered domains. The spectra shown on both panels originate from one of the four phase-separated samples analyzed in this work (same sample as in Fig. S3). For each hydration state and each phase, the spectrum is averaged from at least 10 background-corrected spectra collected from distinct domains within the 20 x 20 µm sample area. The resulting spectra were normalized to better visualize the changes.

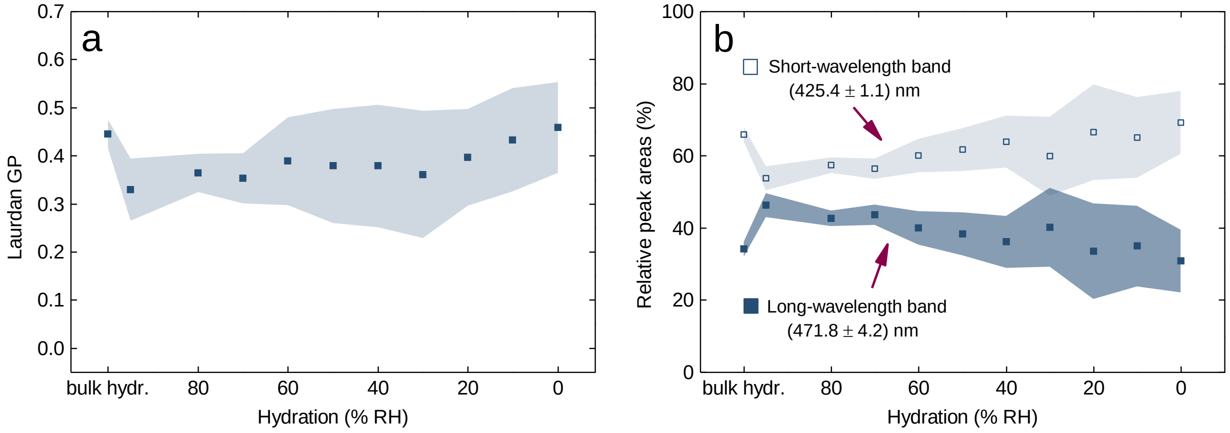


**Figure S5**. (a) Laurdan GP as a function of hydration level of liquid-ordered domains from phase-separated solid supported lipid bilayer composed of an equimolar mixture of di14:1-Δ9*cis*-PC, cholesterol, and egg sphingomyelin. (b) The relative area of the two log-normal functions that give the best fit to the Laurdan emission spectra in the same SLB system as a function of membrane hydration state. Open and full symbols are used for short- and long-wavelength bands, respectively. The data shown on both panels are averaged over four different samples. The uncertainties are standard deviations, denoted as shadows around mean values.

### Note 1.

The insensitivity of Laurdan's emission spectrum in the liquid-ordered phase to changes in hydration over a fairly wide range is unlikely to indicate that no changes are occurring within this phase, but rather that Laurdan is not the appropriate dye to probe them. The liquid-ordered phase composed of saturated phospholipids (egg sphingomyelin in our case) and a high proportion of cholesterol is found to be the least hydrated and the most ordered among all the possible phases.[5,6] It is also reflected in the log-normal decomposition results (Fig. S4b), which revealed the considerably low contribution of long-wavelength band in the Laurdan emission even at fully hydrated conditions compared to liquid-disordered domains or pure phospholipid bilayer. Even if the functional groups of sphingomyelin at the membrane depth where Laurdan resides are partially hydrated, its dipolar relaxation is slow compared to Laurdan's fluorescence timescale.[7] The stiff lipid acyl chains implicate the negligible contribution of Laurdan population surrounded by the relaxable environment to the emission spectrum over a wide range of membrane hydration states. Hence, in the main text we focused on the changes occurring within the liquid-disordered phase, for which the rate of dipolar relaxation is comparable to Laurdan fluorescence lifetime.[7]

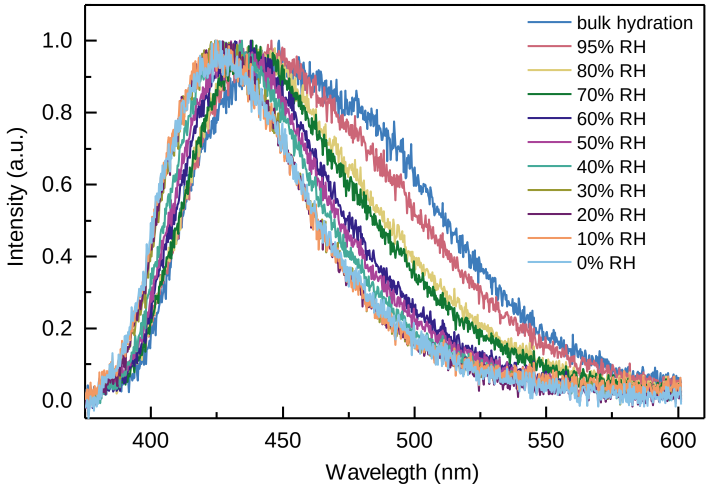


**Figure S6.** Fluorescence spectrum of Laurdan embedded in an exemplary solid-supported lipid bilayer composed of a binary mixture of di14:1-Δ9*cis*-PC and cholesterol at $x_{Chol}$ = 0.3 as a function of membrane hydration state. For each hydration state, the spectrum is averaged from at least 10 background-corrected spectra collected from distinct spots within the 20 x 20 μm sample area. The resulting spectra were normalized to better visualize the changes.

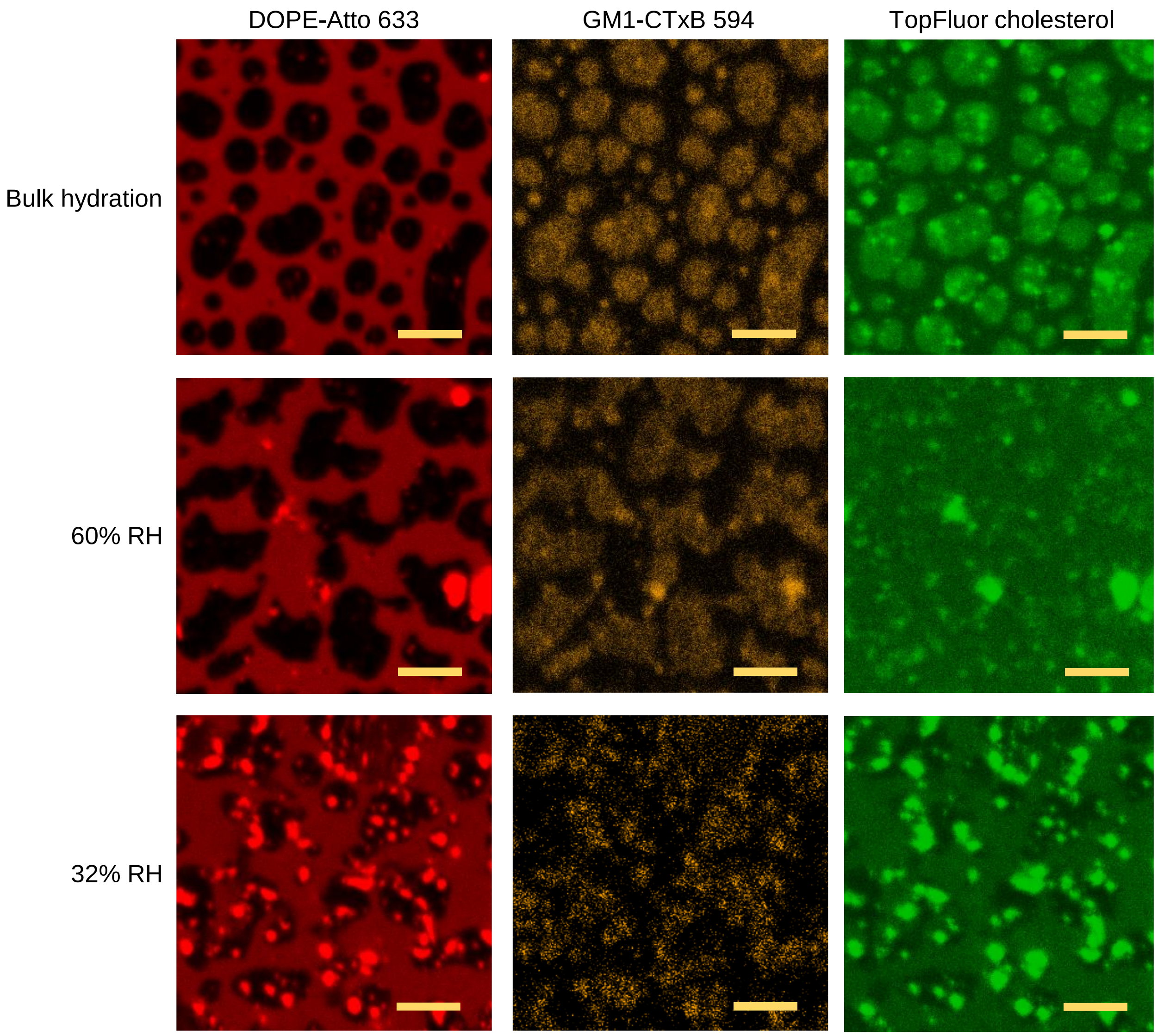


**Figure S7**. Confocal fluorescence microscopy images of a triple-labeled phase-separated solid-supported lipid bilayer composed of an equimolar mixture of di14:1-Δ9*cis*-PC, cholesterol, and egg sphingomyelin for three different membrane hydration states. Liquid-disordered phase is labeled with Atto 633-DOPE (red, left column), liquid-ordered phase is labeled with Alexa Fluor 594-CTxB-GM1 complex (yellow, middle column) and cholesterol, which partitions in both phases is labeled with TopFluor-Chol probe (green, right column). Images for each hydration state originate from different sample area. Under fully hydrated conditions higher intensity of TopFluor-Chol is found in liquid-ordered domains, denoting that liquid-ordered phase contains more cholesterol than liquid-disordered phase, in accordance with the previous reports. At 60% RH the contrast virtually vanishes, indicative of homogenous distribution of cholesterol between distinct phases. At 32% RH the contrast is reversed with respect to the fully hydrated conditions, denoting that at low hydration conditions liquid-disordered phase contains more cholesterol than liquid-ordered phase. The concentration of each dye was 0.1% mol. The scale bar corresponds to 5 μm.
